# ARKS: chromosome-scale scaffolding of human genome drafts with linked read kmers

**DOI:** 10.1101/306902

**Authors:** Lauren Coombe, Jessica Zhang, Benjamin P Vandervalk, Justin Chu, Shaun D Jackman, Inanc Birol, René L Warren

## Abstract

**Background:** The long-range sequencing information captured by linked reads, such as those available from 10x Genomics (10xG), helps resolve genome sequence repeats, and yields accurate and contiguous draft genome assemblies. We introduce ARKS, an alignment-free linked read genome scaffolding methodology that uses linked reads to organize genome assemblies further into contiguous drafts. Our approach departs from other read alignment-dependent linked read scaffolders, including our own (ARCS), and uses a kmer-based mapping approach. The kmer mapping strategy has several advantages over read alignment methods, including better usability and faster processing, as it precludes the need for input sequence formatting and draft sequence assembly indexing. The reliance on kmers instead of read alignments for pairing sequences relaxes the workflow requirements, and drastically reduces the run time.

**Results:** Here, we show how linked reads, when used in conjunction with Hi-C data for scaffolding, improve a draft human genome assembly of PacBio long-read data five-fold (baseline vs. ARKS NG50=4.6 vs. 23.1 Mbp, respectively). We also demonstrate how the method provides further improvements of a megabase-scale Supernova human genome assembly, which itself exclusively uses linked read data for assembly, with an execution speed six to nine times faster than competitive linked read scaffolders. Following ARKS scaffolding of a human genome 10xG Supernova assembly (of cell line NA12878), fewer than 9 scaffolds cover each chromosome, except the largest (chromosome 1, n=13).

**Conclusions:** ARKS uses a kmer mapping strategy instead of linked read alignments to record and associate the barcode information needed to order and orient draft assembly sequences. The simplified workflow, when compared to that of our initial implementation, ARCS, markedly improves run time performances on experimental human genome datasets. Furthermore, ARKS utilizes barcoding information from linked reads to estimate gap size. It accomplishes this by modeling the relationship between known distances of a region within contigs and calculating associated Jaccard indices. ARKS has the potential to provide correct, chromosome-scale, genome assemblies, promptly. We expect ARKS to have broad utility in helping refine draft genomes.

## Background

*De novo* genome assembly, a reference-free approach to reconstructing genome sequences, is essential for establishing reference resources for a molecular-level understanding of living organisms, and for unbiased genomic and genetic analyses in re-sequencing studies. The success of *de novo* genome assembly is highly dependent on the quality and length of the provided sequencing reads. While short sequence reads are typically generated at a higher throughput with greater accuracy and lower cost, the more costly and noisy long reads of current sequencing platforms allow for important resolution of short sequence repeats.

Recently, 10x Genomics (Pleasanton, CA) developed a new library construction technology, which captures long-range information while leveraging the affordability and accuracy of the Illumina sequencing platform (San Diego, CA). The 10x Genomics (10xG) Chromium methodology produces short, barcoded linked reads that co-locate to the same long DNA molecule. This typically provides low sequence coverage (sub 1X) of individual DNA molecules, while generating high molecule coverage genome-wide. It is an effective and less expensive alternative to long reads for phasing nucleotide variants, *de novo* assembly, and linking genome sequences [1–3]. Supernova, 10xG’s own *de novo* assembler, has the ability to output diploid genome assemblies with megabase-scale contiguity from Chromium linked read libraries, and demonstrates the potential presented by this data type for *de novo* genome assembly [2].

Although the most recent, 10xG is not the first to market a linked read technology with the ability to provide long-range sequence information that can be used for ordering and orienting draft genome sequences into scaffolds. Contiguity preserving transposition sequencing data (CPT-seq), which uses the concept of clustering reads from similar genomic regions, is used by the genome scaffolding software fragScaff [4]. In addition, Architect, a scaffolder designed based on Moleculo read clouds specifications, also uses barcoded reads to improve the contiguity of genome assemblies [5]. More recently, we implemented ARCS, the first stand-alone genome scaffolder designed specifically for 10xG linked reads [3]. Although fragScaff and Architect have either demonstrated or suggested that they are applicable to 10xG-derived linked reads, comparisons with ARCS show limitations in their use of this data [3]. All three solutions demonstrate at least some ability to improve draft genomes, but their reliance on read alignments, though effective, negatively impacts their usability and flexibility, as they are dependent on the output of the aligner used and custom formatting requirements of the input linked read sequences.

Although short read alignments are versatile, they are often the most computationally expensive steps in bioinformatics pipelines, and are often avoidable. A growing number of alignment-free bioinformatics software packages have found applications in gene expression quantification [6–8], metagenomic identification and abundance quantification [9–11], and pre-assembly overlap detection for overlap-layout-consensus (OLC) assemblers [12–13]. All have shown increased performance over alignment-based methods.

Here we present ARKS, the Assembly Roundup by linked-read Kmer mapping Scaffolder. ARKS is a streamlined alternative scaffolding methodology that improves upon ARCS by replacing its dependency on read alignments to the draft genome in favor of exact *kmer* matching. This change improves both the software usability and run-time efficiency, with genome scaffolding outcomes comparable to that of ARCS. Read “kmerization” (deconstructing reads into shorter sequences of uniform length *k* using a sliding window) has been explored in countless applications for *de novo* genome assembly, including the ABySS assembler, Supernova assembler and LINKS scaffolder to name a few [2, 14–15]. Methods based on exact *kmer* matching generally eliminate potential alignment software bias, typically have faster processing, and are simpler to execute as the underlying algorithm precludes sequence database indexing and read alignments. Within ARKS, departing from sequence read alignments and moving towards the utilization of *kmers* for pairing contigs effectively eliminates the required upstream steps of input read file formatting and read alignments, which are otherwise tedious when using alignment-based scaffolding software, and vastly improves overall runtime. The faster runtime gives users more control over the adjustment of parameters, which could be challenging when running read alignments with larger (>3 Gbp) genomes. Here, we exhibit ARKS, ARCS, fragScaff, and Architect in three use cases, and show how, using contiguous human genome drafts as baseline assemblies, the ARKS scaffolding algorithm is efficient and effective. ARKS performs so without compromising on the contiguity and accuracy of assembled genomes, yielding results that improve upon the other leading alignment-dependent genome scaffolders assessed.

## Implementation

### ARKS Algorithm

Given a draft genome assembly and a set of linked reads, the goal of ARKS is to produce a scaffold graph in which nodes represent contigs and directed edges represent inferred order of probably adjacent contig pairs within the genome. Briefly, ARKS infers the edges of the graph by determining the Chromium barcodes associated with the 5’-end (head) and 3’-end (tail) sequences of each contig, and connecting all contig head/tail pairs that share more than a minimum number of barcodes. Once ARKS has produced the scaffold graph, downstream scaffolding algorithms such as LINKS can be used to navigate the graph and translate high-confidence paths into scaffold sequences. The principal challenge of the ARKS algorithm is to efficiently identify all probable contig pairs within the draft genome assembly, along with their relative head/tail orientations.

First, for the sake of argument, we make a clear distinction between the terms “alignment” and “mapping”: while the former refers to inferring the relative coordinates of bases between a query and a target, the latter is about determining if the query sequence hits a particular target. The ARKS algorithm proceeds in three steps: (1) mapping Chromium linked barcodes to contigs, (2) scoring candidate contig pairs, and (3) generating the output scaffold graph with estimated distances. Compared to ARCS [3], ARKS uses kmer-based mapping for the first step instead of aligning reads to contigs with BWA mem [16], employs the same algorithm for the second step, and builds on the third step to include distance estimations between neighbouring contigs. In the first step, we map the Chromium reads to the draft assembly sequences (hereinafter inclusively termed contigs referring to contigs and scaffolds for simplicity and clarity) in order to build a set of barcode-to-contig mappings (Fig. 1A), which are used in the second step of the algorithm. More precisely, we build a mapping from each barcode to a list of contig head/tail regions, hereafter referred to as *contig ends*. To reduce memory usage and noise in later steps of the algorithm, we mask the interior sequences of the contigs, and leave only a fixed-length sequence (parameter -*e*, default 30 kbp) at the head and tail of the contig sequence as targets for mapping. To map the reads, we first index the contig ends by shredding the sequences into *kmers*, and by loading them into a hash table that associates *kmers* and contig ends. Then, for each read, we find the best-matching contig end by shredding the read into *kmers*, hashing the *kmers* to identify candidate contig ends, and finally selecting the contig end with the largest fraction of *kmers* overlapping the read using the scoring function

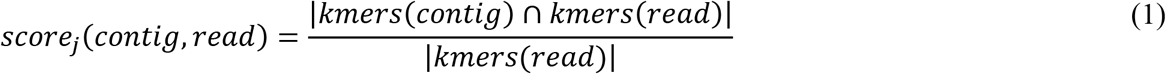

where the operator |·| refers to set cardinality. To reduce spurious mappings, we require a contig end to match a minimum fraction of the read *kmers* (parameter -*j*, 0.55 by default) before it is considered as a candidate.

**Figure 1.**
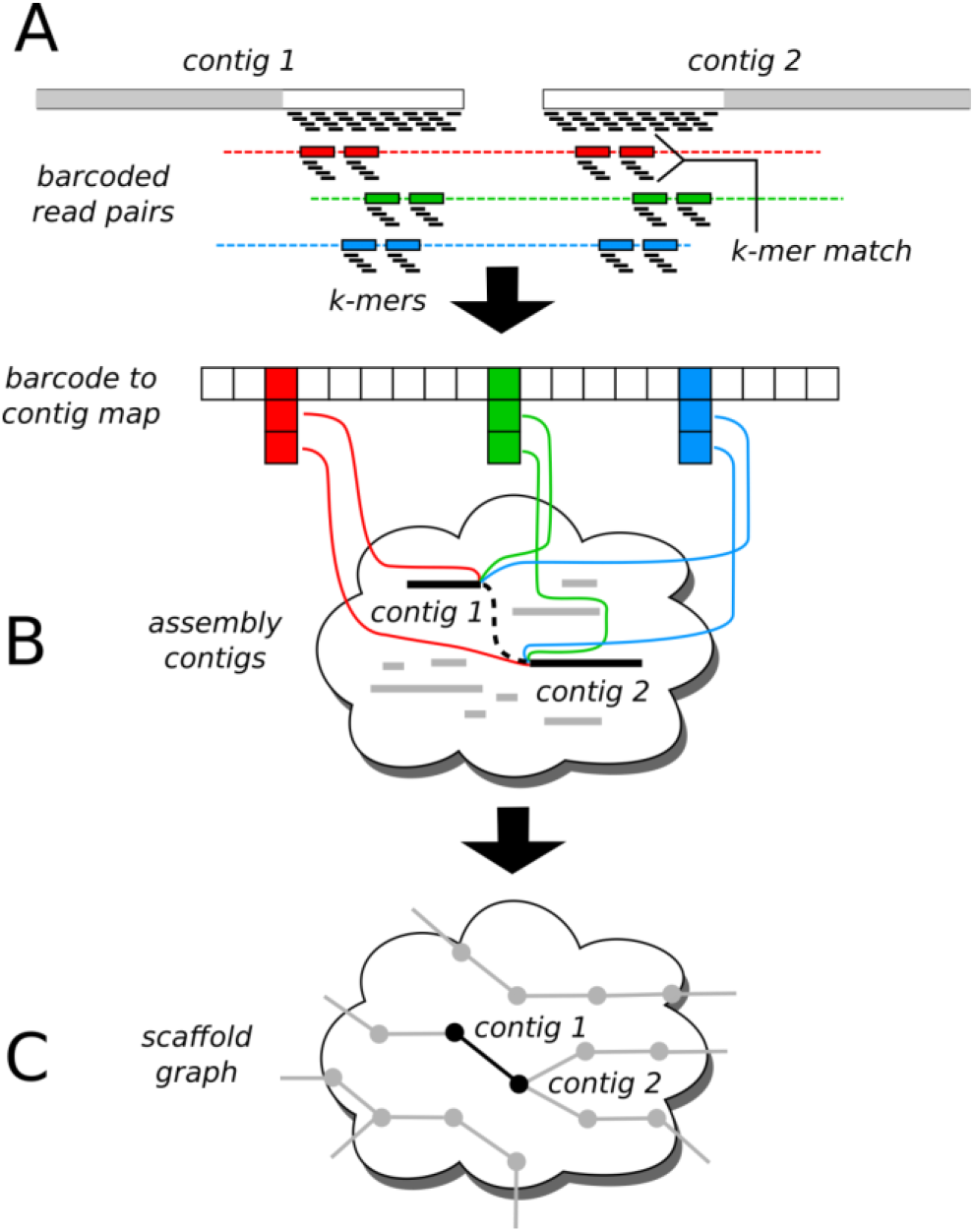
ARKS algorithm. A) In the first step, the barcoded Chromium reads are mapped to the ends of the draft assembly sequences (indicated as contigs in the figure) using a *kmer*-based approach. Reads from three distinct Chromium barcodes are depicted in red, green, and blue, with connecting dashed lines indicating the underlying long DNA molecules for each barcode. The gray regions of the target contigs indicate interior sequence that is masked during mapping. The barcode/contig associations derived from the read mappings are stored in a hash table that maps barcodes to contig ends. (B) In the second step, we iterate over the barcode-to-contig map and tally the number of barcodes that are shared by each candidate pair of contig ends. C) In the third and final step, we generate the output scaffold graph by creating an edge for each candidate pair of contig ends that has greater than a threshold number of shared barcodes (0 by default).

To further increase the specificity of the read mappings, we also require both reads of a pair to map to the same target, and discard *kmers* having multiple contig end memberships. For each read pair that is successfully mapped, we store its barcode in a hash table that maps barcodes to contig ends. This hash table is the input for the second step of the algorithm.

In the second step, we iterate over the hash table we built, in order to identify and score candidate contig end pairs (Fig. 1B). Iterating over the keys (barcodes), we obtain a list of contig ends that match each barcode. Then, for each possible pairing of contig ends within the list, we increment a counter that tracks the number of shared barcodes for the pair. These counts are stored in a second hash table that maps pairs of contig ends to counts, and is used as the input for the third and final step of the algorithm.

In the third step, we generate the edges of the scaffold graph by iterating over the shared barcode counts for each candidate pair of contig ends (Fig. 1C). All pairs of contig ends with a count greater than a minimum threshold (parameter -*l*, 0 by default) are output as an edge of the output graph. It is possible that we will obtain non-zero barcode counts linking the same pair of contigs in multiple orientations, where the possible orientations are head-to-head, head-to-tail, tail-to-head, and tail-to-tail. This can occur, for example, if a long DNA molecule spans the full length of a contig, causing the same barcode to map to both ends. In the case of multiple candidate orientations, we perform a binomial test with the null hypothesis that the two orientations with the highest shared barcode counts are equally likely. If the test generates a *p*-value less than a user-defined threshold (parameter -*r*, 0.05 by default), we output the edge with the orientation corresponding to the highest shared barcode count; otherwise, we omit the edge from the graph.

In generating the scaffold graph, ARKS also uses the barcode data to estimate distances between neighbouring contigs. These constraints are potentially useful for downstream assembly and scaffolding algorithms, in order to identify correct paths. In general, pairs of contig ends that are more distant will share fewer barcodes; however, the parameters of this relationship are dependent on the sequencing coverage and the length distribution of the underlying long molecules. In order to build an empirical model of the barcodes-to-distance relationship, we first train our algorithm using the distances between head and tail regions within given contigs. For each input contig with length greater than two times the head/tail length (parameter -*e* defined above), we record both the distance between the head and tail regions and the Jaccard index for the number of shared barcodes

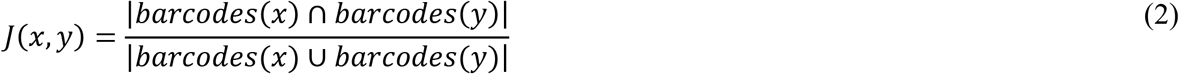

where *x* and *y* are adjacent, linked contig sequences. For a typical empirical relationship between the Jaccard index and the distance, see Additional file 1: Fig. S1. Next, we estimate the distance between the neighbouring contigs by taking the median of intra-contig distance samples with the *B* closest Jaccard indices (parameter -*B*, 20 by default).

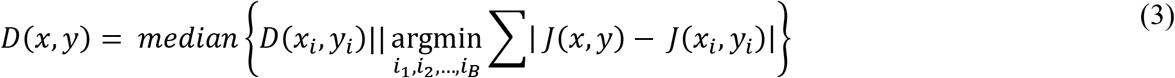

### Data Sources

Human Chromium linked reads were obtained for two individuals’ cell lines, NA24143 and NA12878. The NA24143 reads were downloaded from the Genome in a Bottle (GIAB) FTP site, while the NA12878 reads were downloaded from the 10xG company website (Additional file 1: Table S1). The *C. elegans* Chromium reads were simulated from the Bristol N2 Strain reference using LRSim (v1.0) [17] at 50-fold coverage using parameters -*x* 17 -*f* 50 -*t* 20 -*m* 10, as recommended for genomes of this size range (~100Mbp) (Additional file 1: Table S2). The NA24143 Falcon assembly, as well as its Hi-C/HiRise- scaffolded counterpart, were obtained from the GIAB FTP site (Additional file 1: Table S1).

### Data Analysis

The raw Chromium reads from *C. elegans* as well as the NA12878 human individual reads were processed using Long Ranger (v2.1.3) [18] to extract the barcodes from the reads. The barcodes of the NA24143 reads were already extracted in the downloaded file. For ARCS, fragScaff, and Architect, the Long Ranger output reads required additional read formatting to append the barcode to the read header in the form “header_<barcode>“. Following this formatting step, the reads were aligned to the corresponding draft genome using BWA mem (v0.7.12) [16], setting -*t* 8. ARKS does not require similar read formatting as it reads barcodes directly from the BX tag of the Long Ranger output.

For the *C. elegans* scaffolding runs, the raw simulated 10xG reads were assembled using Supernova (v1.2.0) [2] to produce the draft assembly. Both ARKS and ARCS (v1.0.0) [3] were run with parameters - *c* 8 -*z* 500 -*m* 8-10000 -*e* 30000 and with LINKS (v1.8.5) [15] parameters -*l* 5 and -*a* 0.3, 0.5. A sweep of values for -*k* was tested for ARKS, which was run with eight threads (-*t* 8).

While running fragScaff (v140324) we also used eight threads (-*t* 8), and varied the end node size (parameter -*E*), the mean number of links per bin (parameter -*j*), and the score multiplier (parameter -*u*), as these are suggested to be the most influential scaffolding parameters [4]. We used -*E* 30000 to be consistent with ARKS and ARCS, but also performed runs with -*E* 13508 because it is half of the N80 of the draft assembly, which is approximately the recommended end node size [4]. We used -*j* 1 -*u* 2 to test stringency and improve quality, -*j* 6.5 -*u* 2.5 because Adey *et al*. [4] note that these are the optimal scaffolding parameters for *Drosophila melanogaster*, which has a slightly larger genome than *C. elegans* (175 Mbp vs 100 Mbp), and -*j* 3 -*u* 2 as an intermediate parameter setting.

For the Architect (v0.1) scaffolding runs, the parameters –*rc-abs-thr, –rc-rel-edge-thr*, and –*rc-rel-prun-thr* control the thresholds for creating and pruning a read-cloud edge, respectively, and are suggested to be the most influential parameters [5]. For these *C. elegans* runs, we set –*rc-abs-thr* 5, and vary –*rc-rel-edge-thr* (0.2, 0.3) and –*rc-rel-prun-thr* (0.1, 0.2) to provide a comprehensive sweep.

Raw 10xG Chromium data from the NA12878 individual was assembled using Supernova (v1.2.0) [2]. The draft Supernova assembly was scaffolded using both ARKS and ARCS with parameters -*c* 5 -*e* 30000 -*z* 3000 -*r* 0.05 and -*m* 50-6000. We kept the LINKS parameter -*l* constant at 5, but varied *a*=0.3, 0.5, in decreasing stringency. We ran ARKS with a range of -*k* values (60 to 100, step 20), and with eight threads (-*t* 8). For the fragScaff runs, we set -*E* 30000 to correspond to -*e* in ARKS/ARCS and used eight threads (-*t* 8). We tested -*j* 1 -*u* 2 to ensure high stringency, -*j* 1.75 -*u* 2.5 because Adey *et al*. [4] found these values to be optimal, and -*j* 3 -*u* 2.5 because they were the optimal values for a baseline N50 length of 47 kbp. For the Architect runs, we ran all combinations of –*rc-abs-thr* (10, 5), –*rc-rel-edge-thr* (0.2, 0.3, 0.4), and –*rc-rel-prun-thr* (0.1, 0.2), to provide a comprehensive sweep of parameters.

The NA24143 Falcon and Falcon-HiRise assemblies were corrected using Tigmint [19], which uses Chromium molecule coverage to identify putative misassemblies (parameters *w*=2000, *n*=2). The NA24143 assemblies were then scaffolded using ARKS with parameters -*c* 5, -*g* 1, -*j* 0.5, -*z* 3000, -*e* 30000, -*r* 0.05, -*t* 8, -*m* 50-1000, and LINKS parameter -*l* 5. We parameterized the values of both -*k* and - *a* in distinct scaffolding runs.

To compare the actual distances between merged scaffolds with the distances estimated by ARKS, the *C. elegans* and NA12878 Supernova assemblies were scaffolded with ARKS using the gap distance estimation option (-*D*) (parameters -*c* 8 -*z* 500 -*m* 8-10000 -*e* 30000 -*k* 60 -*a* 0.5 -*j* 0.5 and -*c* 5 -*e* 30000 -*z* 3000 -*m* 50-6000 -*k* 60 -*a* 0.5 -*j* 0.5 for the two genomes, respectively). Actual distances between scaffolds were derived from alignments of the scaffolds to the appropriate reference genomes.

Misassemblies in all genome drafts were assessed using QUAST (v5.0.0, –*scaffolds, –scaffold-gap-max-size* 100000) [20]. All benchmarking was completed using a DELL server with 128 Intel(R) Xeon(R) CPU E7-8867 v3, 2.50GHz with 2.6TB RAM.

## Results and Discussion

We first demonstrate the ability of ARKS to scaffold medium size (~100 Mbp) draft genomes with synthetic *C. elegans* linked read data. We simulated 10xG-like linked reads at 50-fold coverage using LRSim [17] and the reference genome for the Bristol N2 strain (Additional file 1: Tables S1 and S2), and compared the most contiguous assemblies achieved by each tool with QUAST [20]. In comparison to the baseline Supernova assembly, ARKS optimally achieves a genome contiguity, as measured by the NG50 length metric, of 1.11 Mbp, which represents a 4.1-fold increase over the initial baseline draft assembly (Additional file 1: Tables S3 and S4). In comparison, the alignment-dependent ARCS yields a 4-fold increase in contiguity (NG50 = 1.06 Mbp). Architect and fragScaff both yield lower improvements with 1. 5-fold and 3.9-fold increases in assembly contiguity, respectively. While the assemblies generated by ARCS and ARKS have a similar number of misassemblies (207 and 217, respectively), the number of reported misassemblies for the fragScaff assembly is roughly quadruple, with 959. Compared to the ARKS and ARCS assemblies, the fragScaff assembly has a higher number of sequence inversions (6 vs. 5 vs. 31, respectively), and translocations (56 vs. 55 vs. 86, respectively). Both ARCS and ARKS offer similar contiguity and accuracy on the simulated *C. elegans* dataset, and represent an improvement over alternative scaffolding methodologies assessed. Finally, the distance estimates from ARKS strongly correlate with the actual distances between merged scaffolds elucidated from the Bristol N2 strain reference genome, with the estimated and actual distances having a Pearson correlation coefficient of 0. 896 and a mean absolute error of 2.9 kbp (Additional file 1: Fig. S2A).

Next, we used ARKS to further scaffold an already contiguous human genome assembly (NG50 = 14.7 Mbp) generated with Supernova (Additional file 1: Table S3). The same high-depth (~156-fold) Chromium reads from the NA12878 individual used to generate the baseline Supernova assembly were subsequently used for scaffolding (Additional file 1: Table S2). ARKS increased the baseline NG50 length 1.8-fold (25.94 Mbp compared to 14.74 Mbp for the baseline), demonstrating that ARKS is able to make additional sequence merges while using the same linked read information as was provided to Supernova (Fig. 2A). In contrast, ARCS and fragScaff increased the baseline NG50 length 1.6 and 1.3- fold in this order (NG50=23.09 and 19.06 Mbp, respectively), while Architect did not produce significant improvements over the parameters tested. Although less stringent fragScaff parameters generated an assembly comparable to the contiguity obtained by ARCS and ARKS (NG50 = 23.37 Mbp), this was at the expense of additional misassemblies (2889 for fragScaff vs. 1441 for ARKS; Additional file 1: Table S5). When comparing the accuracy of the most contiguous ARCS and ARKS assemblies, we observe that both harbor a very similar number of misassemblies (1473 and 1441, respectively), and they represent a 11. 9% and 9.5% increase over the number of misassemblies identified in the baseline Supernova assembly.

**Figure 2.**
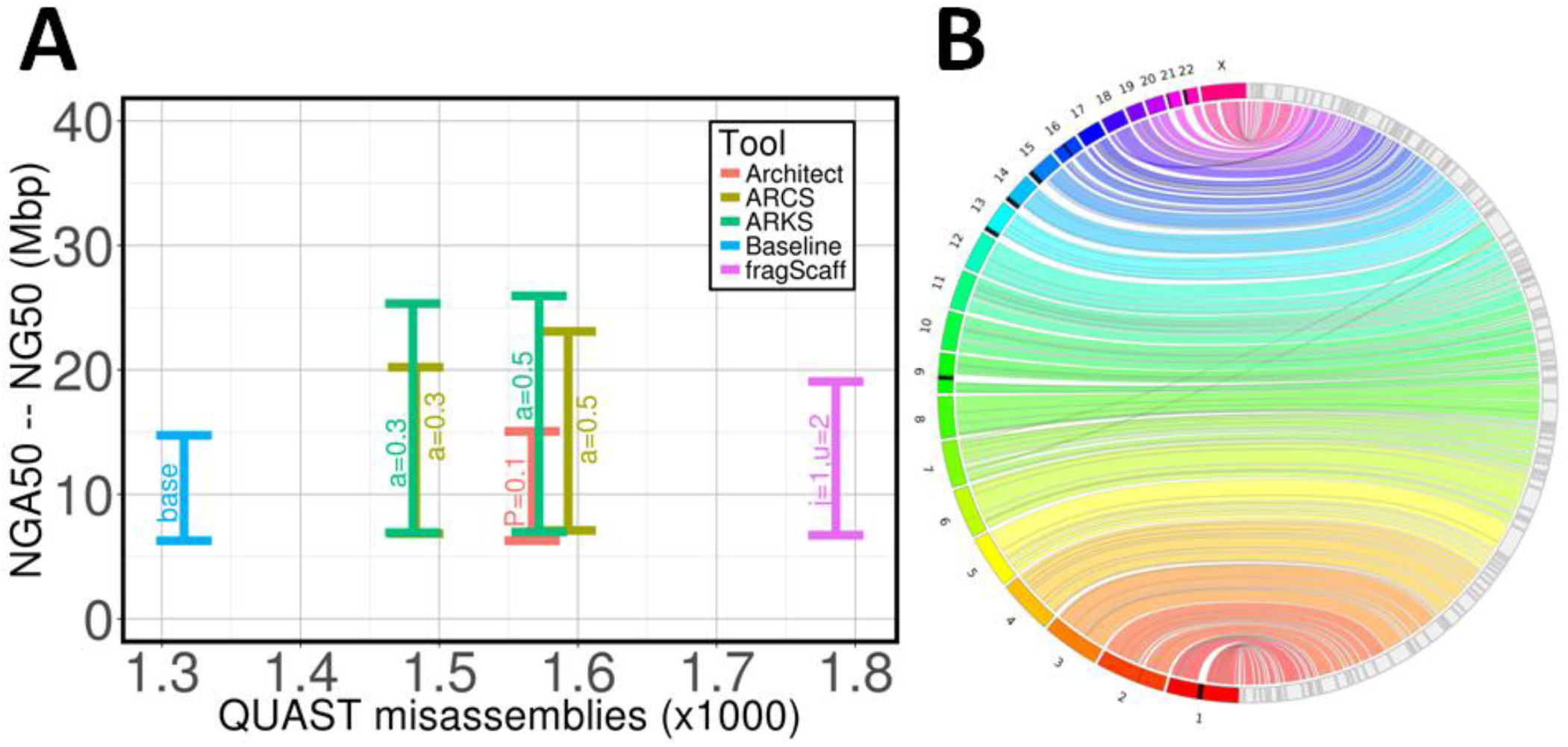
Scaffolding a 10xG Supernova human genome assembly with ARKS. A) Comparing the contiguity and accuracy of assemblies scaffolded by ARkS, ARCS, fragScaff and Architect as measured by QUAST. The baseline NA12878 Supernova assembly was scaffolded using ARCS (-c5 -s98 -m50-6000 -z3000 -e30000), ARKS (-c5 -k100 -t8 -j0.5 -m50-6000 -z3000 -e30000), fragScaff (-E30000 -j1 -u2) and Architect (–rc-abs-thr5 –rc-rel-edge-thr0.2), –rc-rel-prun-thr abbreviated to ‘P’). The Y-axes show the range of NGA50 to NG50 lengths to indicate the uncertainty caused by real genomic variations between individual NA24143 and the reference genome GRCh38. B) A Circos [24] assembly consistency plot of ARKS (-k100 -j0.5 -c5 -e30000 -z3000 -m50-6000 -r0.05 -a0.5) scaffolding of the baseline NA12878 Supernova assembly. Scaftigs from the largest 123 scaffolds, consisting of 85% (NG85) of the genome, are aligned to GRCh38 with BWA mem [16]. GRCh38 chromosomes are displayed incrementally from 1 (bottom, red) to X (top, fuchsia) on the left while scaffolds (grey with black outlines) are displayed on the right side of the rim. Connections show aligned regions, 1 Mbp and larger, between the genome and scaffolds. Large-scale misassemblies are visible as cross-over ribbons. The black regions on chromosomes indicate reconstruction gaps in the reference. The majority of each chromosome is represented in the final ARKS assembly by no more than 13 assembly scaffolds.

Comparing the largest (NG85) ARKS scaffolds to the reference human genome GRCh38, we do not detect any large structural rearrangements nor inversions (Fig. 2B). The majority (22/23) of the human chromosomes in the genome are represented in the ARKS assembly by fewer than nine scaffolds (Fig. 2B, Fig. 3, Additional file 1: Table S6). Furthermore, for most of the human chromosomes (21/23), baseline NG85 scaffolds are merged by ARKS, thus noticeably improving the chromosome-scale contiguity of the Supernova baseline assembly (Fig. 3). Notably, chromosomes 20 and 21 are represented by two scaffolds each - the number expected due to the presence of centromeres. In both cases, the two scaffolds reconstruct most of the chromosome’s bases (93.9% for chromosome 20, and 97.2% for chromosome 21; Additional file 1: Table S6; Fig. 2B; Fig. 3). Note that there are few discrepancies between the number of NG85 scaffolds visible in Figure 3 and listed in Additional file 1 (Table S6) due to constituent contigs of a scaffold (ie. scaftigs) aligning non-collinearly. These cases could be caused by misassemblies or structural variation in the NA12878 cell line compared to the GRCh38 reference.

**Figure 3.**
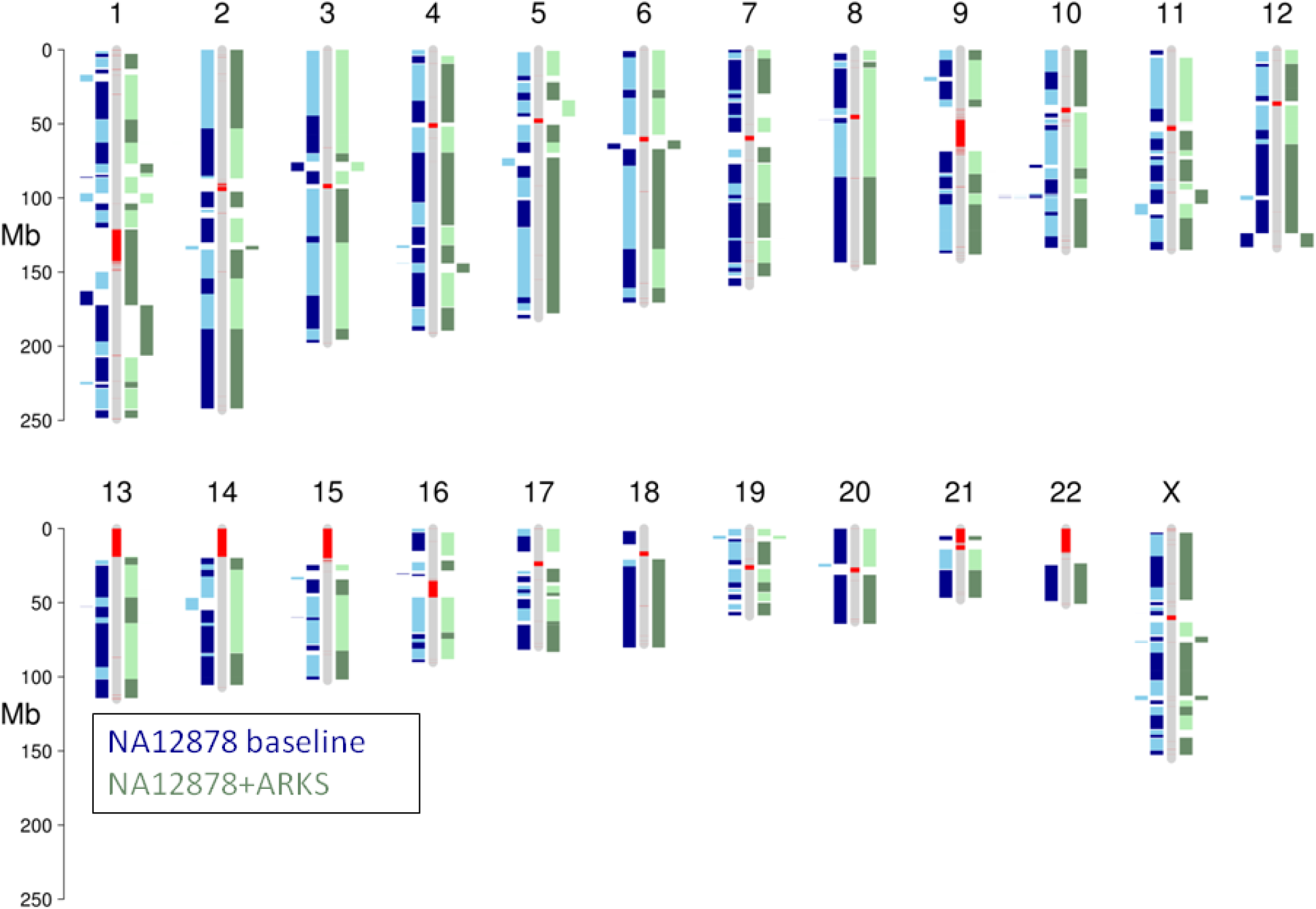
Chromosome-scale ARKS scaffolding of a NA12878 10xG Supernova assembly. Scaftigs from the NG85 scaffolds of the baseline NA12878 Supernova assembly (blue) and the assembly following ARKS scaffolding (-k100 -j0.5 -c5 - e30000 -z3000 -m50-6000 -r0.05 -a0.5) (green) were aligned to GRCh38 using BWA mem [16]. The ideogram was generated using the R package chromPlot [25]. For each assembly, alternating shades represent different scaffolds. Red bands on the reference chromosomes denote gaps in the reference assembly.

When scaffolding the NA12878 Supernova assembly with ARKS using the distance estimation feature, the distance estimates produced by ARKS strongly correlate with the true distances between merged scaffolds, with a Pearson correlation coefficient of 0.872 and a mean absolute error of 8.6 kb (Additional file 1: Fig. S2B). Thus, the ARKS distance estimates provide useful information about the gap sizes between merged contigs. In addition to showcasing the utility of the distance estimation functionality in ARKS, the measures of the actual distances between sequences also demonstrate the ability of ARKS to leverage the long-range information afforded by the 10xG molecules to scaffold over tens of kilobases, with over 150 merges made between sequences separated by at least 10 kbp.

Time and memory usage for ARCS, fragScaff, and Architect are primarily bottlenecked by the read alignment step preceding the contig pairing/scaffolding stage. Although read alignments can be partitioned into sets, and each set into threads to speed up the alignment step, this is also at the expense of additional read formatting and higher computational resource requirements. Because ARKS records the location of every *kmer* found, ARKS’s main bottleneck is memory, and is caused by kmerization of contig ends. We note that a more fragmented draft will have higher memory requirements since many more contig ends will need to be shredded into *kmers* (Additional file 1: Tables S7 and S8). Notwithstanding, our results show that ARKS is still comparable to ARCS, fragScaff, and Architect in memory requirements using the sequence assembly data tested here (Additional file 1: Tables S9, S10 and S11). In terms of run time, the kmerization step within ARKS effectively eliminates the need for input read formatting and alignments, and shortens the overall wall clock time for scaffolding the NA12878 Supernova assembly almost six-fold (10.5h vs. 58.9h vs. 71.3h vs. 96.9h for ARKS, ARCS, fragScaff and Architect, respectively) (Fig. 4, Additional file 1: Table S10).

**Figure 4.**
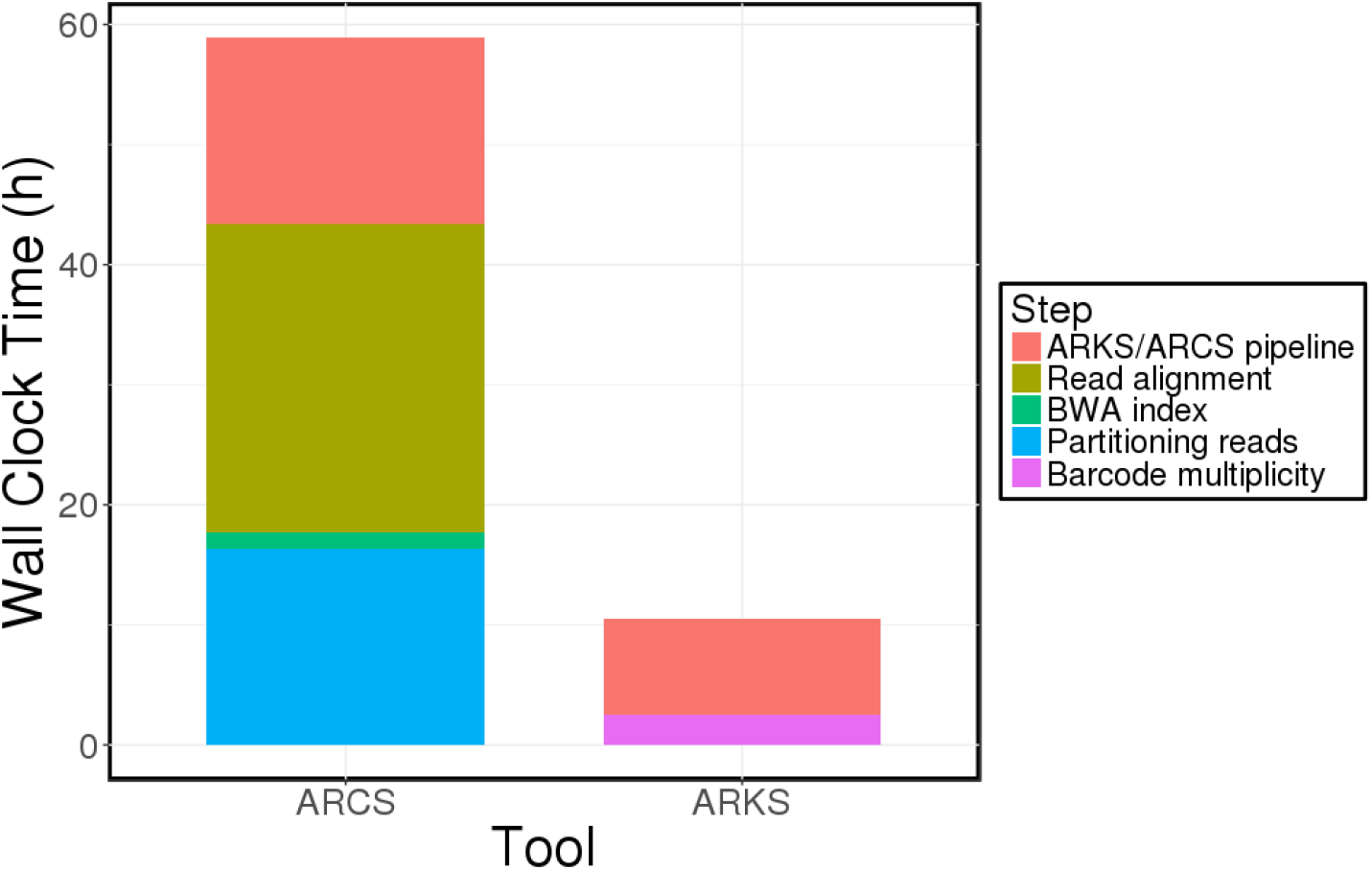
Benchmarking wall clock time for ARCS and ARKS. The wall clock benchmarking is shown for the most contiguous assemblies from scaffolding the NA12878 Supernova assembly with ARCS (-a0.5 -s98 -c5 -m50-6000 -e30000 - z3000) and ARKS (-a0.5 -k100 -t8 -c5 -j0.5 -m50-6000 -e30000 -z3000). For ARCS, the linked reads were partitioned into eight sets, and aligned to the draft assembly in parallel. The wall clock time for the bottleneck read alignment is shown. ARKS was run with eight threads, while only the alignment step of ARCS was run with eight threads, as threading is not implemented in ARCS.

In addition to using ARKS to improve a Supernova assembly, we demonstrate the utility of our tool in scaffolding a contiguous human genome (NA24143) draft generated by assembling PacBio reads with Falcon (NG50 = 4.56 Mbp; Additional file 1: Table S3) [21]. Scaffolding the NA24143 Falcon assembly using linked reads with ARKS rivals the scaffolding improvements achieved using Hi-C/HiRise (Dovetail Genomics Inc.), with the scaffolders improving the NG50 length of the assembly 3.3-fold and 3.2-fold, respectively (15.0 Mbp and 14.5 Mbp compared to 4.56 Mbp for the baseline) (Fig. 5; Additional file 1: Table S12). Furthermore, the NG50 length increases 3.7-fold when the NA24143 Falcon assembly is corrected with misassembly corrector Tigmint [19] prior to ARKS scaffolding (NG50 = 16.7 Mbp). In addition to increased contiguity, the assembly generated by running Tigmint followed by ARKS has fewer misassemblies than both the baseline NA24143 Falcon assembly and the Falcon + Hi-C/HiRise assembly (3053 vs. 3100 vs. 3477) (Fig. 5; Additional file 1: Table S12). This demonstrates that scaffolding using 10xG linked reads may be used instead of or in addition to HiC/HiRise scaffolding to generate more contiguous and correct drafts.

**Figure 5.**
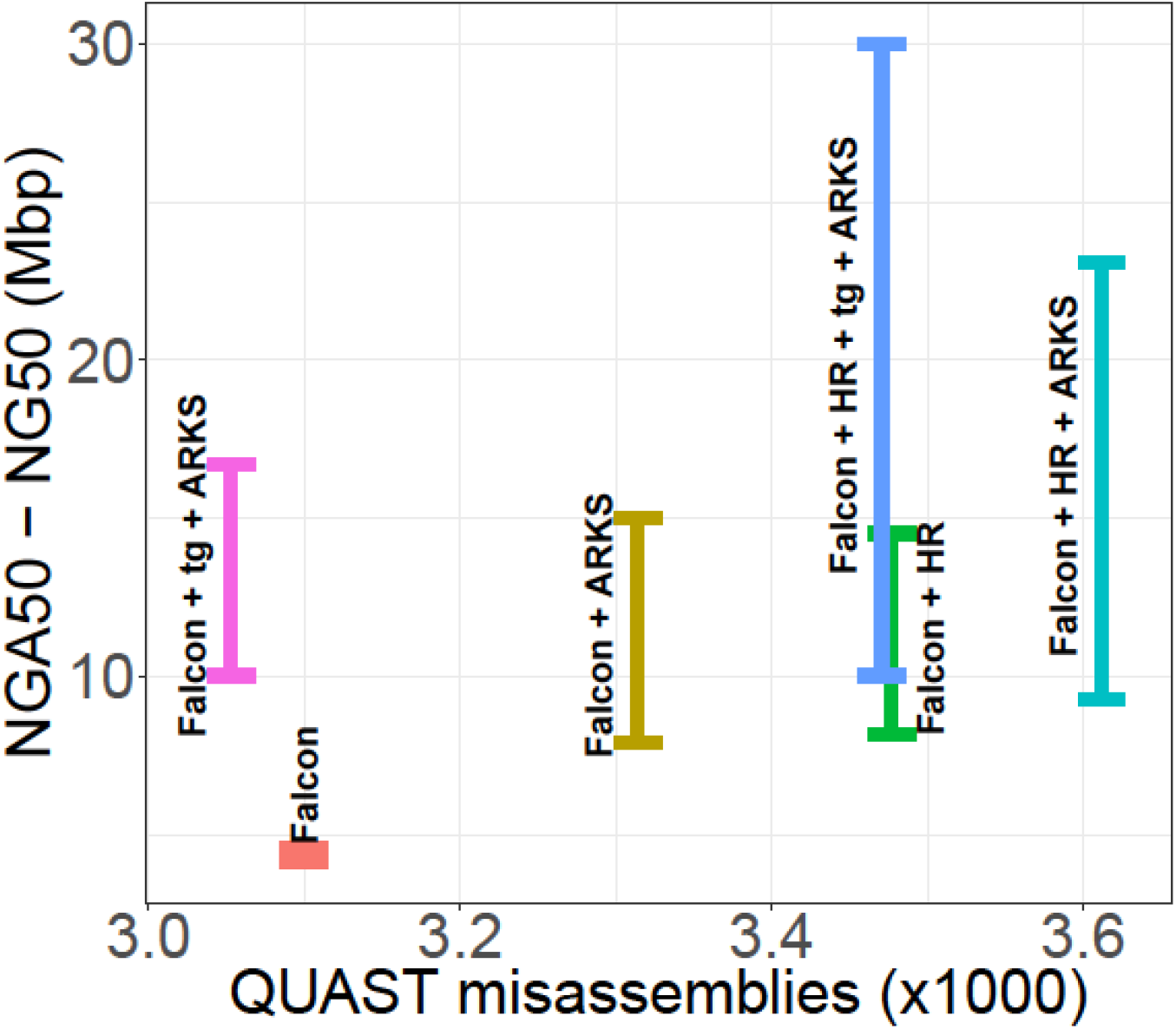
Contiguity gains from scaffolding a NA24143 PacBio Falcon + HiRise human genome assembly with ARKS. The base Falcon (orange bar) and Falcon + HiRise (HR) (green) draft genomes were corrected with Tigmint (tg) [19] (w=2000, n=2), then scaffolded using ARKS (-a0.3 -c5 -k40 -j0.5 -e30000 -z3000 -t8 -m50-1000) (pink and blue bars). We also ran ARKS on the base Falcon (yellow) and Falcon+HiRise (turquoise) draft genomes without Tigmint correction.

Perhaps even more striking is the compounding effect of dual scaffolding, effectively using 10xG/ARKS in conjunction with Hi-C/HiRise. We have previously reported that ARCS works optimally with already contiguous genome drafts as input (>1Mbp N50 length), and this is also a characteristic of ARKS [3]. ARKS scaffolding of the PacBio/Falcon + Hi-C/HiRise scaffolded draft increases the NG50 length of the resulting assembly draft 1.6-fold (23.09 Mbp compared to 14.53 Mbp), and represents a more than 5-fold increase in contiguity over the PacBio/Falcon baseline assembly (23.09 compared to 4.56 Mbp for the baseline) (Fig. 5). Similar to the PacBio/Falcon assembly scaffolding results, the contiguity gains are even greater when running Tigmint [19] prior to ARKS, with a 6.6-fold and 2.1-fold increase in NG50 length over the PacBio/Falcon and PacBio/Falcon + HiC/HiRise assemblies, respectively (30.01 Mbp compared to 4.56 Mbp and 14.53 Mbp, respectively). Furthermore, the additional ARKS scaffolding step only increased the number of misassemblies by 3.9%, while the misassemblies decreased by 0.2% when Tigmint was run prior to ARKS, demonstrating that the sequence merges made by ARKS significantly increased the contiguity of the draft assembly without impeding its accuracy (Fig. 5; Additional file 1: Table S12).

## Conclusions

Further assembly of individual human genome drafts that used the native, 10xG *de novo* assembler Supernova [2] highlights opportunities in retrospectively improving pre-existing and contiguous, megabase range assemblies with linked reads. Our results of further scaffolding Supernova assemblies demonstrates that additional sequence merges are possible even though the same linked read data is reutilized for scaffolding. This will greatly benefit genome-sequencing projects done exclusively using the 10xG Chromium platform, since no additional sequencing or costly scaffolding-specific data is needed and chromosome-scale contiguity is readily achievable. For projects that employ both linked read and Hi-C structural mapping data, such as that presented here with the PacBio long-read assemblies of NA24143 [22–23], we stress that superior chromosome-scale scaffolding is attained when using ARKS in conjunction with the HiRise scaffolder. With ARKS, we show that capturing additional long-range information for scaffolding is possible without read alignments and with better overall resource efficiency, usability and maximum return in contiguity, without compromising on the resulting accuracy of the assembly.

## Availability and requirements

**Project name:** ARKS

**Project home page:** https://github.com/bcgsc/arks

**Operating system:** Platform independent

**Programming language:** C++

**Other requirements:** LINKS (v1.8.6 or higher), Boost, GCC

**License:** GNU GPL

**Any restrictions to use by non-academics:** No

**List of abbreviations**
10xG: 10x Genomics
GIAB: Genome in a Bottle

## Declarations

### Funding

This work has been partly supported by the National Human Genome Research Institute of the National Institutes of Health (under award number R01HG007182). Additional funds were received through Genome Canada and Genome British Columbia (project BCB11101) as well as Genome Canada, Genome Quebec, Genome British Columbia and Genome Alberta for the Spruce-Up (243FOR) project (www.spruce-up.ca). The content reported here is solely the responsibility of the authors, and does not necessarily represent the official views of the National Institutes of Health or other funding organizations.

### Availability of data and material

ARKS is freely available at https://github.com/bcgsc/arks.

Sources for all data used and/or analysed during the current study are listed in Additional file 1: Table S1.

### Authors’ contributions

RLW, IB and JC designed the research; JZ, BPV and SDJ developed the software. LC and JZ ran benchmarking, LC and RW analyzed the data, and LC, RW, JC and BPV made the figures. LC, JZ, RLW, BPV and SDJ wrote the manuscript and all authors contributed to editing of the manuscript. All authors read and approved the final manuscript.

### Ethics approval and consent to participate

Not applicable.

### Consent for publication

Not applicable.

### Competing interests

The authors declare that they have no competing interests.

## Acknowledgments

Not applicable.

## Additional files

Additional file 1: Supplementary Tables and Figures. (PDF 426 KB)

